# Analysis of environmental data during a lagrangian experiment: The influence of vertical movements

**DOI:** 10.1101/2025.10.23.680922

**Authors:** Patricia Romero Fernández, Eulogio Vargas-García, Ricardo Borrego-Santos, Raúl Laiz-Carrión, José Quintanilla, Michael R. Landry, Sven A. Kranz, Michael R. Stukel, Thomas B. Kelly, Manuel Vargas-Yáñez

## Abstract

Investigating the time evolution of physical and biochemical properties of the ocean with *in situ* sampling can follow two approaches: Eulerian and Lagrangian. In the Eulerian approach, repeated measurements are taken at fixed locations, whereas the sampling point moves with the displacement of a water parcel in the Lagrangian approach. During the BLOOFINZ-IO cruise off northwest Australia, four Lagrangian experiments (“cycles”) were conducted with multidisciplinary sampling done at regular intervals for several days following a satellite tracked drifter with mixed-layer drogue. To test the Lagrangian nature of these experiments, we adapted the Bindoff and McDougall (1994) approach for decomposing observed changes between adjacent CTD profiles into components due to vertical movement (heaving) and those occurring along isopycnal surfaces. Profile depth variability was mainly driven by vertical displacements of isopycnals (internal waves), while temperature, chlorophyll, oxygen and salinity were relatively stable when observed on isopycnal surfaces across all casts within the same cycle, and different on average between cycles. Our analysis clearly indicated that density surface was a more appropriate vertical coordinate than physical depth for assessing real environmental variability during each cycle and confirmed the Lagrangian character of the experiments. While the BLOOFINZ cruise serves as a case study, the methodology can be readily extended to analyze other biochemical variables in different ocean regions.

## 1. Introduction

Tuna species play a crucial role in global fisheries, with particularly significant economic implications for certain coastal nations (FAO, 2024; ISSF, 2022). Beyond their commercial value, these species are vital for maintaining the health of marine ecosystems. As apex predators, they exert top-down control over trophic food webs (Estes et al., 2011; Bornatowski et al., 2017), making their sustainable management essential. Achieving this, requires a comprehensive understanding of the natural variability in tuna populations, which is largely driven by annual recruitment success. This success is especially sensitive to larval-stage survival (Niebla et al., 2014; Quintanilla et al., 2024).

Numerous factors influence larval survival. Environmental variables such as temperature can accelerate growth during early developmental stages (Wexler et al., 2011; Kim et al., 2015). Additionally, maternal inheritance, trophic position and niche occupied by larvae, shaped by feeding success, prey availability and interspecific competition, are also critical determinants (Laíz-Carrión et al., 2019; Gerard et al., 2022; Quintanilla et al., 2023). Given the complexity and interconnectedness of these influences, an end-to-end process analysis can be productive, as exemplified by the study of larvae of Atlantic Bluefin Tuna (ABT; *Thunnus thynnus*) in the Gulf of Mexico by the BLOOFINZ-GoM (Bluefin Larvae in Oligotrophic Ocean Foodwebs: Investigations of Nutrients to Zooplankton) Project (Gerard et al., 2022). That integrative approach included analyses of hydrography, nutrient concentrations, primary productivity, larval growth, zooplankton grazing, and isotopic composition to infer trophic levels and interpret larval feeding and growth success.

Conventional Eulerian approaches, which rely on fixed oceanographic stations, may fail to capture the dynamic processes affecting larval survival. Stukel et al. (In review) suggest that observed variations in plankton communities in such experiments are more likely driven by physical processes, such as advection or diffusion, than by biological interactions. Kehinde et al. (2023), Kelly et al. (2021) evidenced the importance of both vertical and lateral advection, especially in oligotrophic waters as those of the Argo basin. This underscores the value of the Lagrangian approach, which tracks water parcels or larval patches through space and time, providing more precise representations of ecosystem dynamics (Satoh et al., 2008).

Gerard et al. (2022) applied this methodology, conducting five multi-day Lagrangian experiments in their study of ABT larvae in the Gulf of Mexico. Their end-to-end process approach revealed key aspects of larval life history, including feeding behavior, dietary selectivity, maternal effects and trophic position that significantly advanced understanding of the environmental and biological conditions influencing larval development under both optimal and deficient conditions (Malca et al., 2022; Shiroza et al., 2022; Quintanilla et al., 2023, 2024).

In 2022, the BLOOFINZ-IO project used a similar approach to conduct a comparative end-to-end habitat study for larval Southern Bluefin Tuna (SBT, *Thunnus maccoyii*) in oligotrophic waters overlying the Argo Abyssal Plain (hereafter Argo Basin) in the eastern Indian Ocean, their only known global spawning ground. The study was carried out during the January–February peak spawning season (Davis et al., 1990; Jenkins and Davis, 1990), and the experimental design comprised four Lagrangian experiments, each involving several days of sampling following a satellite-tracked drift array. In concept, this approach enabled the continuous monitoring of the same water mass, and thus the same larval patch, throughout each cycle.

Here, we test whether these experiments conform to the Lagrangian concept of consistently sampling water with the same characteristics throughout each cycle experiment. Following the approach of Bindoff and McDougall (1994), we detail the methodology for decomposing observed depth variability in cycle profiles at fixed depth/pressure into: (1) a heaving component (vertical displacement, internal waves) versus (2) changes along isopycnal surfaces. We also present the statistical framework for comparing environmental conditions across cycles after vertical displacements have been accounted for.

## 2. Materials and methods

### 2.1. Lagrangian cycles

Cruise RR2201 of the BLOOFINZ Program took place aboard R/V *Roger Revelle* from 26 January to 11 March 2022. As part of the study, four multi-day Lagrangian experiments (hereafter, called “cycles”) were conducted between 3 and 22 February in the southern Argo Basin (Fig. 1A, B). Each cycle involved the deployment of a drifter array with a surface buoy, a 3-m holey-sock drogue centered at 15-m depth, and attachment points on the line, under the float, for *in situ* bottle incubations in net bags (Fig. 1C). The vessel tracked each drifter for several days, conducting repeated CTD casts, water column sampling, and oblique plankton tows at stations spaced several hours apart. Table 1 summarizes the duration, start and end positions, and number of stations for each cycle. Tables S1-S4, in supplementary material, provide detailed information on CTD casts, coordinates, and timestamps for all cycles.

**Figure 1.**
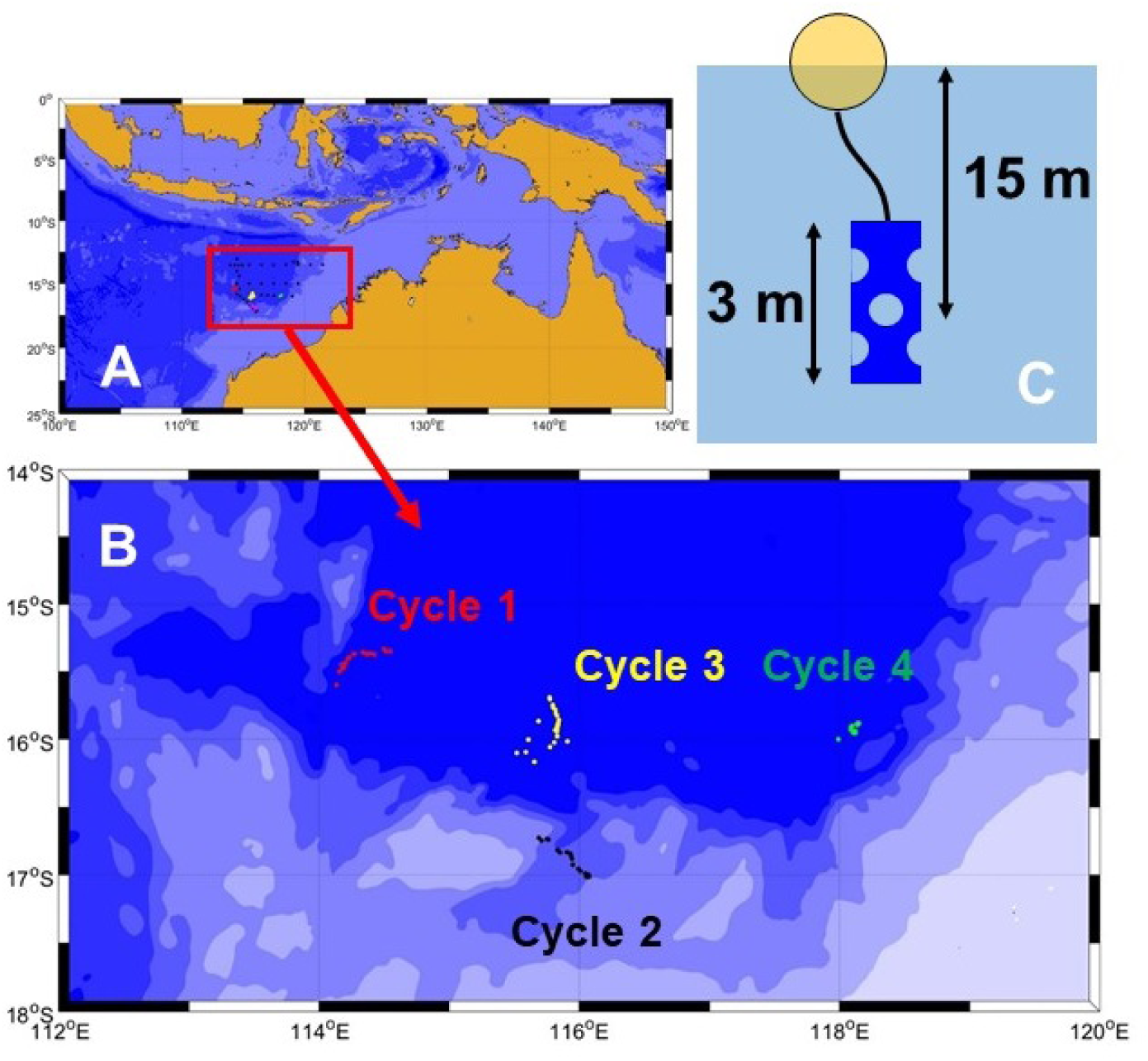
Locations of Lagrangian cycle experiments conducted in the southern Argo Basin during BLOOFINZ cruise RR2201. A, B are regional and expanded maps of CTD cast positions for Cycle 1 (red), Cycle 2 (black), Cycle 3 (yellow) and Cycle 4 (green). C is schematic to the drifter float and drogue.

**Table 1.** Initial and final dates, coordinates and number of CTD casts for each of the 4 Lagrangian experiments conducted on cruise RR2201.

| Cycle | Initial date | Initial coordinates | Final Date | Final coordinates | No. of casts |
| --- | --- | --- | --- | --- | --- |
| 1 | 3 February | 15.35°S, 114.55°E | 7 February | 15.60°S, 114.14°E | 27 |
| 2 | 9 February | 16.73°S, 115.69°E | 12 February | 17.01°S, 116.08°E | 23 |
| 3 | 14 February | 16.10°S, 115.59°E | 18 February | 15.69°S, 115.77°E | 30 |
| 4 | 19 February | 16.00°S, 117.99°E | 22 February | 15.94°S, 118.12°E | 19 |

The CTD was equipped with sensors delivering continuous 1-m resolved vertical profiles of temperature, salinity, density, chlorophyll fluorescence and dissolved oxygen at each station. Note that calibration of CTD data is required using dissolved oxygen concentrations from water samples (Winkler titration; Strickland & Parsons, 1972), and chlorophyll concentrations measured by spectrophotometry or fluorometry (Holm-Hansen et al., 1965; Jeffrey and Humphrey, 1975). However, as described in the next subsection, this study focuses exclusively on the relative variation of these variables. Thus, the CTD profiles were adequate for our purposes in the present work.

### 2.2. Meteorological data

*R/V Roger Revelle* was equipped with a meteorological station that acquired air temperature, atmospheric pressure, precipitation rates, short- and long-wave radiation, and relative wind speed and direction. Navigation data allowed us to calculate absolute wind speed and direction. Besides this, the vessel had an underway seawater system with temperature, conductivity (for salinity calculation), dissolved oxygen, and fluorescence (chlorophyll-a concentration) sensors. Meteorological and sea surface data were acquired at 15-s intervals. These data were averaged to produce a time series with a one-hour resolution. As most meteorological variables are affected by a strong daily cycle, these time series were also smoothed using a 25-point mean average.

### 2.3. Decomposition of observed changes by depth layer

Bindoff and McDougall (1994) analyzed changes in temperature and salinity profiles collected from the same geographic region but measured years apart, aiming to understand the impacts of climate change. They demonstrated that an observed increase (or decrease) in temperature at a given depth did not necessarily indicate warming (or cooling) of the water mass present at that depth. Nor could such changes be directly linked to increased (or decreased) heat absorption at the ocean surface during the formation of that water mass. Similarly, changes in salinity at a specific depth could not be directly attributed to variations in freshwater input at the surface.

Equation (1), adapted from Bindoff and McDougall’s original formulation, replaces potential temperature with a generic variable *φ*, which may represent chlorophyll, dissolved oxygen, temperature, salinity, etc. The equation shows that changes observed at a fixed pressure level (*p*) can be decomposed into two components: one due to changes along isopycnals (*σ*) and another due to vertical displacement of those isopycnals (heaving).

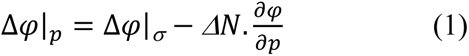

According to Bindoff and McDougall, the left-hand side (LHS) of Equation (1) represents the observed change in variable *φ* at pressure level *p* between two consecutive profiles, where pressure is the vertical coordinate. In the present study, instead of comparing profiles taken years apart, we analyze consecutive CTD casts within the same Lagrangian cycle. Here, *Δ* denotes change. The first term on the right-hand side (RHS) corresponds to changes at constant sigma-theta (*σ*), while *ΔN* represents the vertical displacement of the isopycnals. The final term, *∂φ/∂p*, is the vertical gradient of *φ*. Combined, -*ΔN · ∂φ/∂p* quantifies the contribution of vertical motion (heaving) to the observed change in *φ*.

The decomposition is illustrated in Figure 2. For the sake of clarity, we used simulated chlorophyll profiles in this schematic explanation. Figure 2A shows two chlorophyll profiles from a single Lagrangian experiment, with several sigma-theta surfaces plotted in blue. In this simulation, isopycnals between 50 and 120 dbar were displaced vertically, with a maximum upward shift of 20 dbar occurring at 85 dbar. Subtracting the first profile from the second gives the change in chlorophyll at each depth or pressure level (i.e., the LHS of Equation 1). For example, at 65 dbar, the chlorophyll concentration increased from 0.17 to 0.83 mg/m^3^, a net change of 0.66 mg/ m^3^. Since both profiles were collected while tracking the same water mass, horizontal advection can be initially ruled out, and the change can be attributed to biological processes (e.g., net primary production, photo-acclimation, etc.).

**Figure 2.**
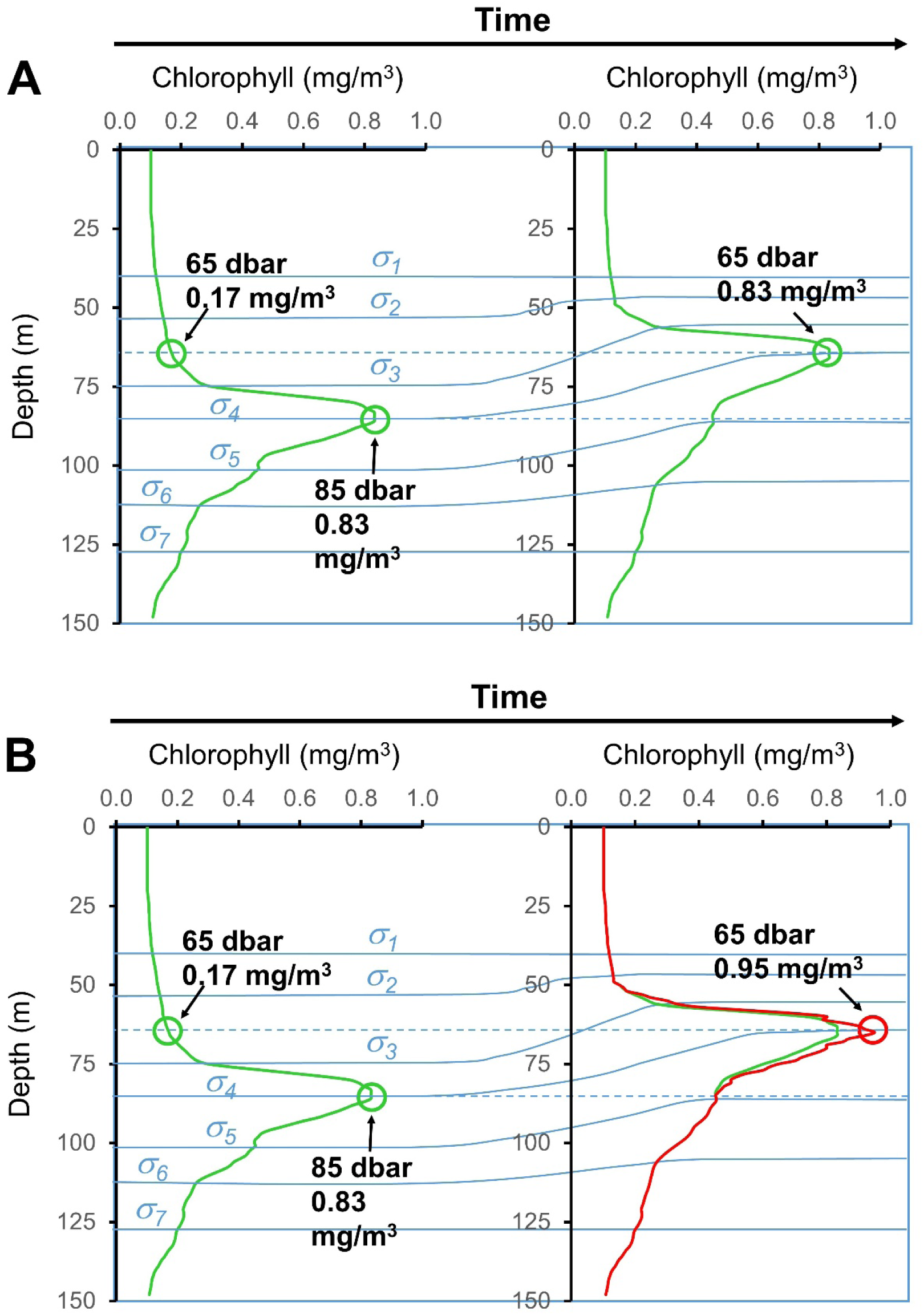
Simulated chlorophyll profiles corresponding to two consecutive CTD casts in which there is an upward movement of isopycnals. A. Interpretation for no change of chlorophyll at constant density. B. Interpretation for a change in chlorophyll (red) relative to no change of chlorophyll at constant density (green).

However, if we account for vertical movement, we see that the *σ₄* isopycnal was located at 85 dbar during the first profile and had risen to 65 dbar in the second. The same water parcel would thus appear higher in the water column, contributing 0.66 mg/ m^3^ to the increase — a change due to heaving, represented by the second RHS term in Equation (1). Now consider a different case illustrated in Figure 2B: the second cast yields the red profile, and isopycnals undergo the same vertical displacement as in Figure 2A. However, this time, the chlorophyll concentration at the *σ₄*isopycnal changes from 0.83 to 0.95 mg/m^3^, an increase of 0.12 mg/ m^3^, which is the first term on the RHS of Equation (1). Since we are following the same water parcel both horizontally (with the drifter) and vertically (along the isopycnal), this increase can be attributed to biological activity. Thus, the total observed increase of 0.78 mg/m^3^ at 65 dbar can be decomposed into 0.12 mg/m^3^ from biological processes along the isopycnal and 0.66 mg/m^3^ from vertical displacement (heaving).

The rationale behind this decomposition is twofold. First, vertical displacements occurring over short timescales, on the order of a few hours as in the cycles analyzed in this study, are very likely driven by internal tides. These tidal motions cause upward and downward movements of water masses that introduce noise into the vertical profiles of various biogeochemical variables. Therefore, if environmental variables were used to study biological processes or to assess their potential influence on larval tuna growth, the heaving contribution should be removed to avoid misleading interpretations.

Second, large differences in physical or biological variables between consecutive profiles could raise doubts about whether the sampling truly followed a Lagrangian path. Each cycle was designed to follow a drifter centered at 15 m, which typically lies within the surface mixed layer where tuna larvae are found (Davis et al., 1990). Thus, it is reasonable to assume that this mixed layer moves as a coherent unit. However, if this assumption fails, due to vertical shear in ocean currents, it is possible that water samples collected while following the drifter may not actually represent the same water parcel or the same phyto- and zooplankton communities.

It is important to note that this analysis assumes that density behaves as a conservative property over the short time scales considered for each cycle. As a result, any observed change in density at a fixed depth is interpreted as a vertical displacement, allowing us to quantify the contribution of such displacements to the variability of other variables, such as chlorophyll or oxygen concentration. However, surface processes like precipitation or heat exchange can modify salinity and/or temperature at the ocean surface.

A decrease in surface density caused by freshwater input or surface warming does not compromise the method, as the resulting density values typically do not match any values from the previous profile, thereby preventing the decomposition. In such cases, the corresponding analyses at these surface levels were omitted. Conversely, surface cooling or intense evaporation can increase surface density and potentially trigger vertical mixing. In such scenarios, the method used here may incorrectly interpret mixing as an upward displacement of isopycnals.

To minimize this source of error, the upper 5 dbar of the water column were excluded from the analysis. Additionally, a 5-point moving average was applied to smooth the vertical density profiles, effectively extending the excluded region to the upper 7 dbar. This potential limitation of the method is further considered in the discussion.

### 2.4. Comparisons of cycle profiles

For each cycle and each variable analyzed in this study (temperature, salinity, sigma-theta, chlorophyll, and oxygen concentration), we obtained a corresponding set of vertical profiles. When our analyses confirmed that all profiles within a given cycle originated from the same water parcel and exhibited consistent physical and biogeochemical conditions, we summarized these conditions using a mean profile. This approach enabled comparisons of mean profiles across different cycles, such as evaluating differences in average temperature profiles among the four cycles.

However, mean profiles may resemble each other in some depth or density layers while differing in others. To account for this variability, comparisons were conducted at specific pressure or density levels. For clarity, consider the example of chlorophyll concentration. At each pressure level, we collected 27 values from Cycle 1, 23 values from Cycle 2, 30 values from Cycle 3, and 19 values from Cycle 4, each corresponding to the number of casts per cycle. For instance, chlorophyll concentrations measured at 25 dbar in each cycle were treated as samples from four independent distributions with known means and standard deviations. The objective was to test whether these means differed significantly across the cycles.

Shapiro-Wilk tests were initially applied to evaluate the normality of the data at each pressure level (e.g., 25 dbar). If normality was confirmed, an ANOVA was used to test for differences in means among the cycles. If normality was not met, we used the non-parametric Kruskal-Wallis test (H-statistic) instead. This analysis was repeated across all pressure levels and for all variables, recording the resulting F-statistics (ANOVA) or H-statistics (Kruskal-Wallis), their critical values (α = 0.05), and the associated p-values. The same methodology was applied using sigma-theta as the vertical coordinate.

## 3. Results

### 3.1. Chlorophyll and oxygen profiles from paired casts

Changes in chlorophyll and dissolved oxygen concentrations were calculated for each pair of consecutive casts within the same cycle. These changes were initially computed at constant pressure levels and subsequently decomposed into two components: variations along isopycnal surfaces and changes induced by heaving (vertical displacement). Figure 3 illustrates these results. Figure 3A shows profiles for chlorophyll (green lines) and dissolved oxygen (blue lines) for cast 11 (solid lines) and cast 12 (dashed lines) from Cycle 1. Figures 3B (green line) and 3C (blue line) display the differences between the two casts—i.e., the changes in chlorophyll and oxygen concentrations at fixed pressure levels. In both Figures 3B, C, the black lines represent the contribution from heaving, while the red lines indicate changes occurring along isopycnal surfaces. The dotted green line in Figure 3B and the dotted blue line in Figure 3C represent the sum of both components for chlorophyll and oxygen, respectively, which correspond, as expected, to the total changes observed at constant pressure levels. Figures 3B, C show that the heaving contributions are much larger than the changes on isopycnal surfaces. As an additional example, Figures 3D-F show the same comparison of changes on isopycnals surfaces versus heaving for Cycle 1 casts 13 and 14.

**Figure 3.**
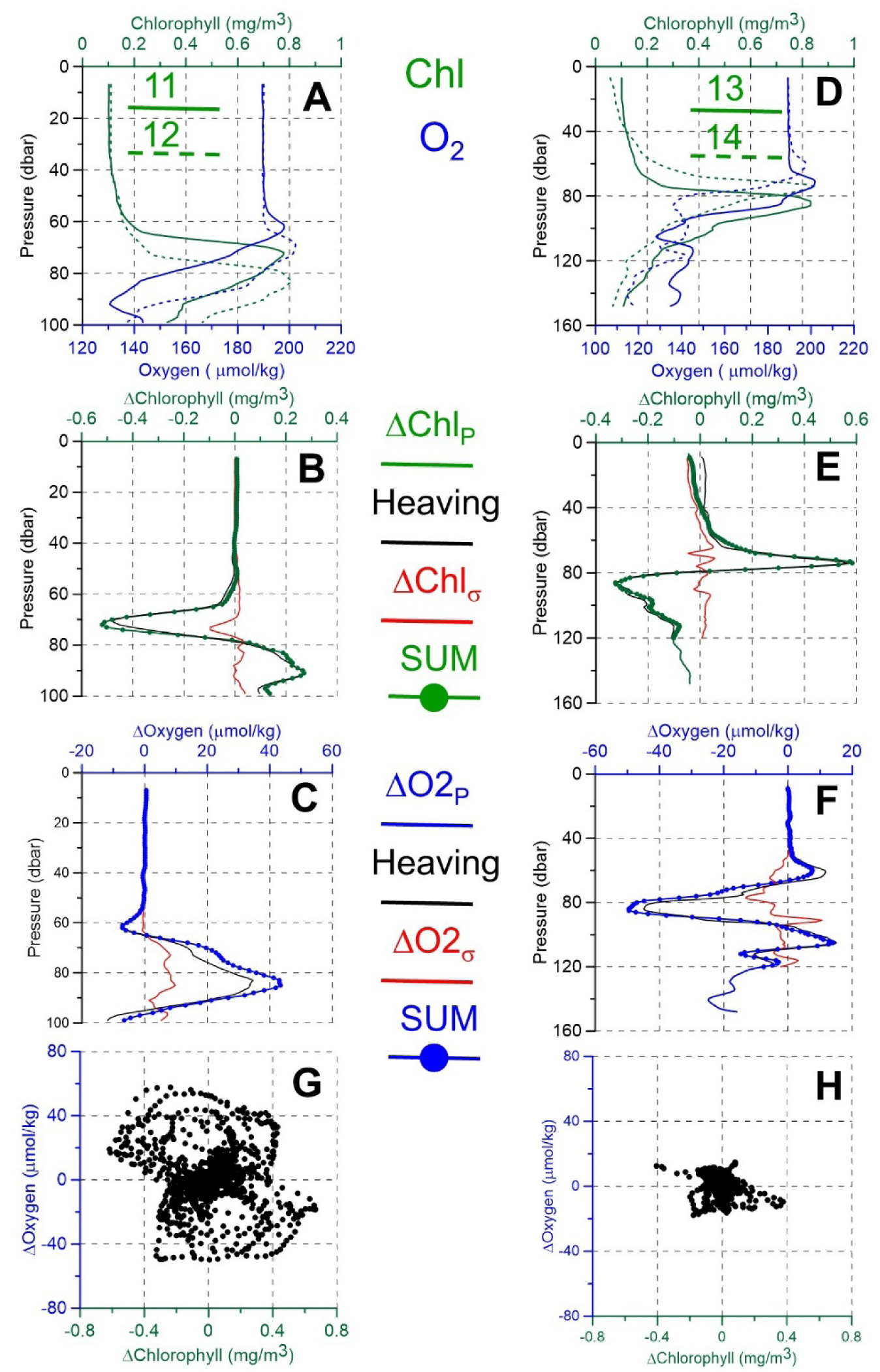
Changes in chlorophyll and dissolved oxygen concentrations between Cycle 1 casts 11 and 12 (panels A-C) and between Cycle 1 casts 13 and 14 (panels D-F). Panel A shows chlorophyll (green lines) and dissolved oxygen (blue lines) profiles for casts 11 (solid lines) and 12 (dashed lines). Panel D shows chlorophyll (green lines) and dissolved oxygen (blue lines) profiles for casts 13 (solid lines) and 14 (dashed lines). Panels B, E show changes at constant pressure for chlorophyll (green line); red line is change along isopycnals; black line is heaving; green dotted line is sum of heaving and isopycnal contributions. Panels C, F are the same decomposition as B, E but for dissolved oxygen concentration (blue line); heaving (black line), isopycnal changes (red lines). Panel G shows scatter plot of changes in dissolved oxygen against chlorophyll for all pairs of Cycle 1 consecutive casts and pressure levels. Panel H shows scatter plot of changes in dissolved oxygen against chlorophyll calculated on isopycnal surfaces.

This analysis was repeated for all pairs of casts across the four cycles. Figure 3G presents changes in dissolved oxygen versus changes in chlorophyll concentration at constant pressure across all levels (surface to 150 dbar) and all cast pairs for Cycle 1. Figure 3H shows the same relationship, but calculated along isopycnal surfaces with identical axis ranges to enable a direct visual comparison to Figure 3G. The results indicate that while changes at constant pressure can reach up to ±0.8 mg m^-3^ (chlorophyll) and ±60 µmol kg^-1^ (dissolved oxygen), these variations are substantially reduced when evaluated along isopycnal surfaces. This confirms that heaving is the primary driver of the large magnitude of observed changes.

### 3.2. Temporal evolution of water mass properties

In order to have a clearer picture of the evolution of water mass properties throughout the four cycles, Figure 4 shows Hovmöller diagrams for sigma-theta, chlorophyll and oxygen concentration for each of the cycles. For the clarity of the plot, cast numbers have been used instead of time for the x-axes (including days, hours and minutes was too cluttered). However, and in order to have an idea of the time of the day when the profiles were obtained, Figure S1 in supplementary material shows the temporal evolution of chlorophyll concentration using sigma-theta as vertical coordinates including symbols to indicate the time of midnight and noon throughout each cycle (see also discussion section). Figures 4A-E correspond to Cycle 1, whereas panels 4F-J, 4K-O and 4P-T correspond to Cycles 2, 3 and 4, respectively. In all the cases, the first of the plot panels (for instance, Figure 4A for Cycle 1) presents the evolution of sigma-theta, showing clearly the vertical displacements of isopycnals. The following two panels (for instance, B and C for Cycle 1) show the evolution of chlorophyll concentration. In the first case, (Figure 4B), the vertical coordinate is pressure, and chlorophyll exhibits large fluctuations that resemble those observed for the isopycnal surfaces (Fig. 4A). On the contrary, chlorophyll isolines flatten considerably when sigma-theta is used as vertical coordinate, reflecting that vertical displacements constitute a large fraction of the chlorophyll variability. The same behavior was observed for dissolved oxygen concentrations (panels 4D-E for Cycle 1, 4I-J for Cycle 2, 4N-O for Cycle 3 and 4S-T for Cycle 4). In order to avoid extrapolation, shaded areas correspond to those sigma-theta values not found for a certain chlorophyll or dissolved oxygen profile. This can be the case when the most surface sigma-theta in a certain profile is higher than the first sigma-theta level or simply when the profile did not reach the bottom or the highest sigma-theta value.

**Figure 4.**
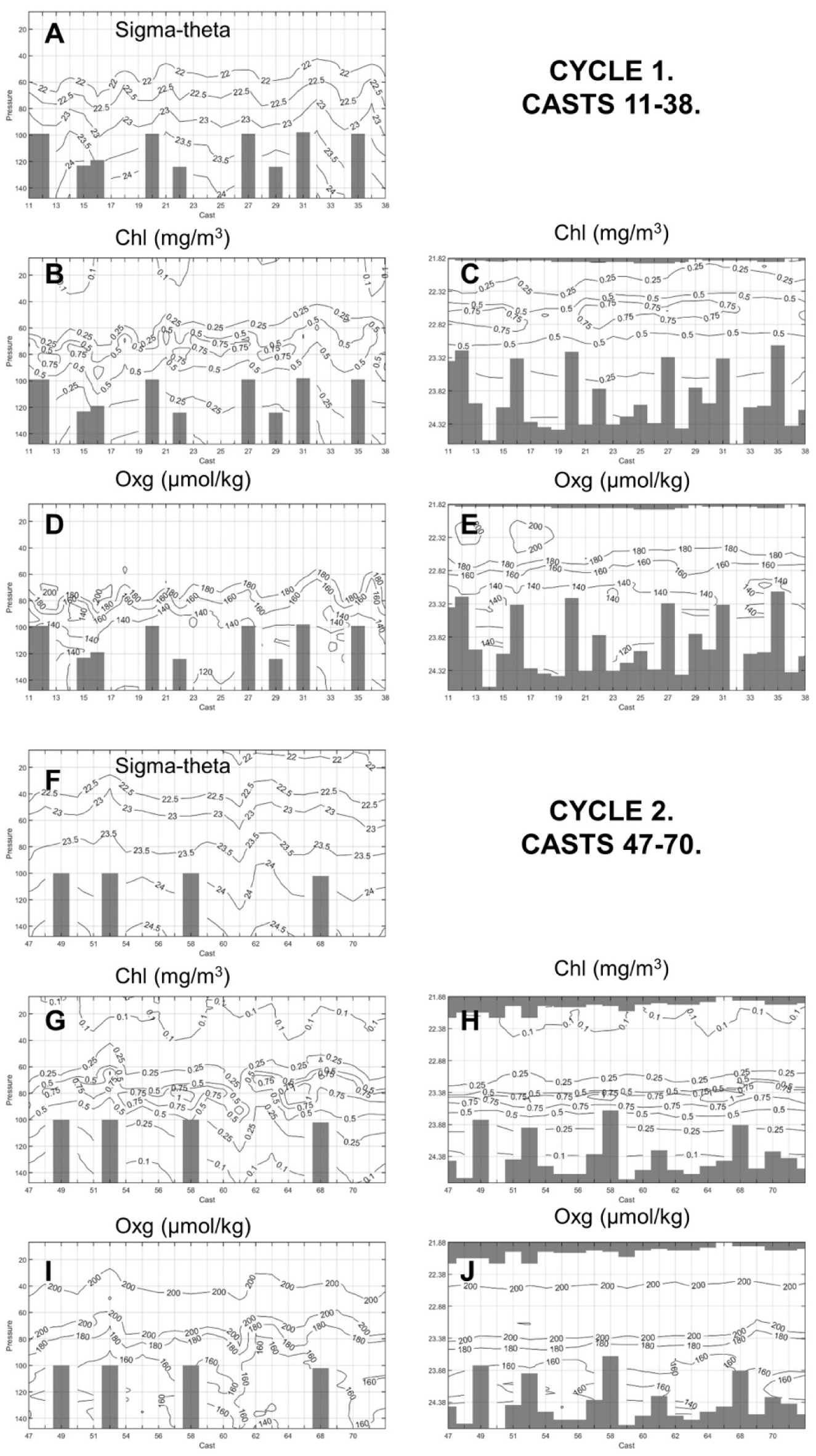
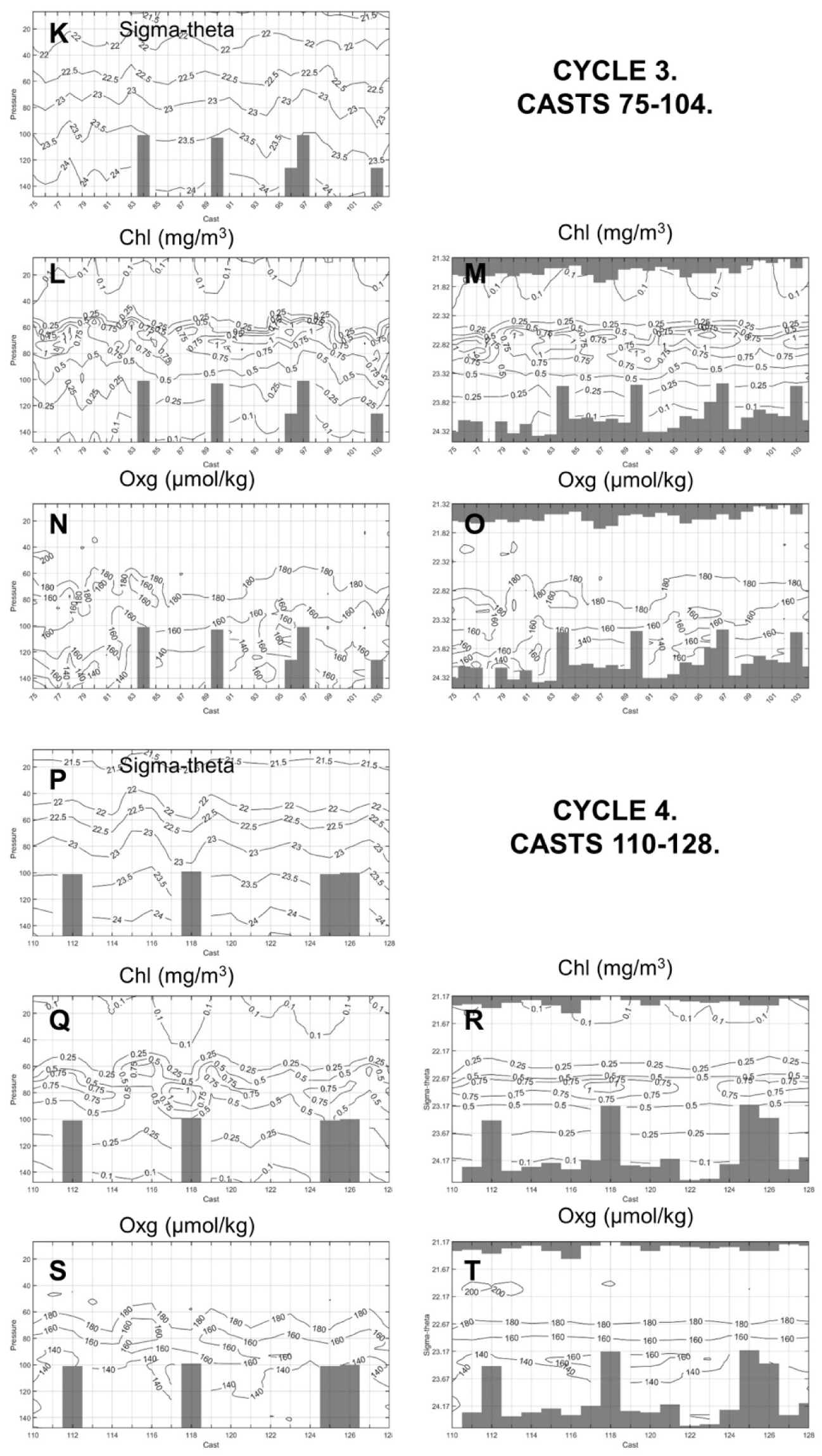
Figures A, F, K and P show the time evolution (Hovmöller diagrams) for the vertical sigma-theta profiles for Cycles 1 to 4: Figures 4B,C (Cycle 1), 4G,H (Cycle 2), 4L,M (Cycle 3) and 4Q,R (Cycle 4) show the time evolution of chlorophyll concentration using the pressure and sigma-theta as vertical coordinates respectively. Figures 4D,E (Cycle 1), 4I,J (Cycle 2), 4N,O (Cycle 3) and 4S,T (Cycle 4) show the time evolution of dissolved oxygen concentration using the pressure and sigma-theta as vertical coordinates respectively. Sigma-theta is expressed in kg/m^3^ -1000, chlorophyll concentration in mg/m3 and dissolved oxygen in µmol/kg. Gray bars are unsampled depths on individual casts.

When sigma-theta is considered as the vertical coordinate (i.e. when we analyze the evolution of chlorophyll and dissolved oxygen on isopycnal surfaces), profiles obtained from the same cycle (notably Cycles 1, 2 and 4) are very similar, indicating that, in fact, they correspond to the same water parcel. However, Cycle 3 shows a larger variability and could be considered as an exception to this general behavior.

The same analysis was conducted for temperature and salinity (see Figure S2 in supplementary material). It is interesting to note that temperature profiles are very stable within each cycle when sigma-theta is used as vertical coordinate. Figures S2 B, F, J and N show that, if we consider the changes of temperature on isopycnal surfaces, the same water mass was being sampled throughout the complete cycle. If the pressure was considered as vertical coordinate, that is, changes were calculated at fixed depth or pressure levels, some changes could be appreciated (Supplementary Figure S2). However, these results show that such changes were simply caused by vertical displacements of material surfaces.

### 3.3. Profile variability based on isobaric versus isopycnal surfaces

The variability of temperature, salinity, chlorophyll and oxygen concentration depend on whether these variables were observed at depth levels (isobaric levels) or on isopycnal surfaces. Dots in Figures 5A,B show all the chlorophyll vertical profiles corresponding to the four cycles. Solid lines show the mean profiles averaged for each cycle. In Figure 5A, the pressure is the vertical coordinate, whereas in Figure 5B, it is sigma-theta. Figures 5 C,D show the same comparison but for temperature. In order to quantify the differences in Figure 5, standard deviations were calculated for each cycle and for each pressure or density level. These results are presented in Figure 6. Figures 6 A,B show the standard deviations of chlorophyll concentration for each cycle as a function of pressure (Fig. 6A) and versus sigma-theta (Fig. 6B). Figures 6 C,D show similar results for temperature.

**Figure 5.**
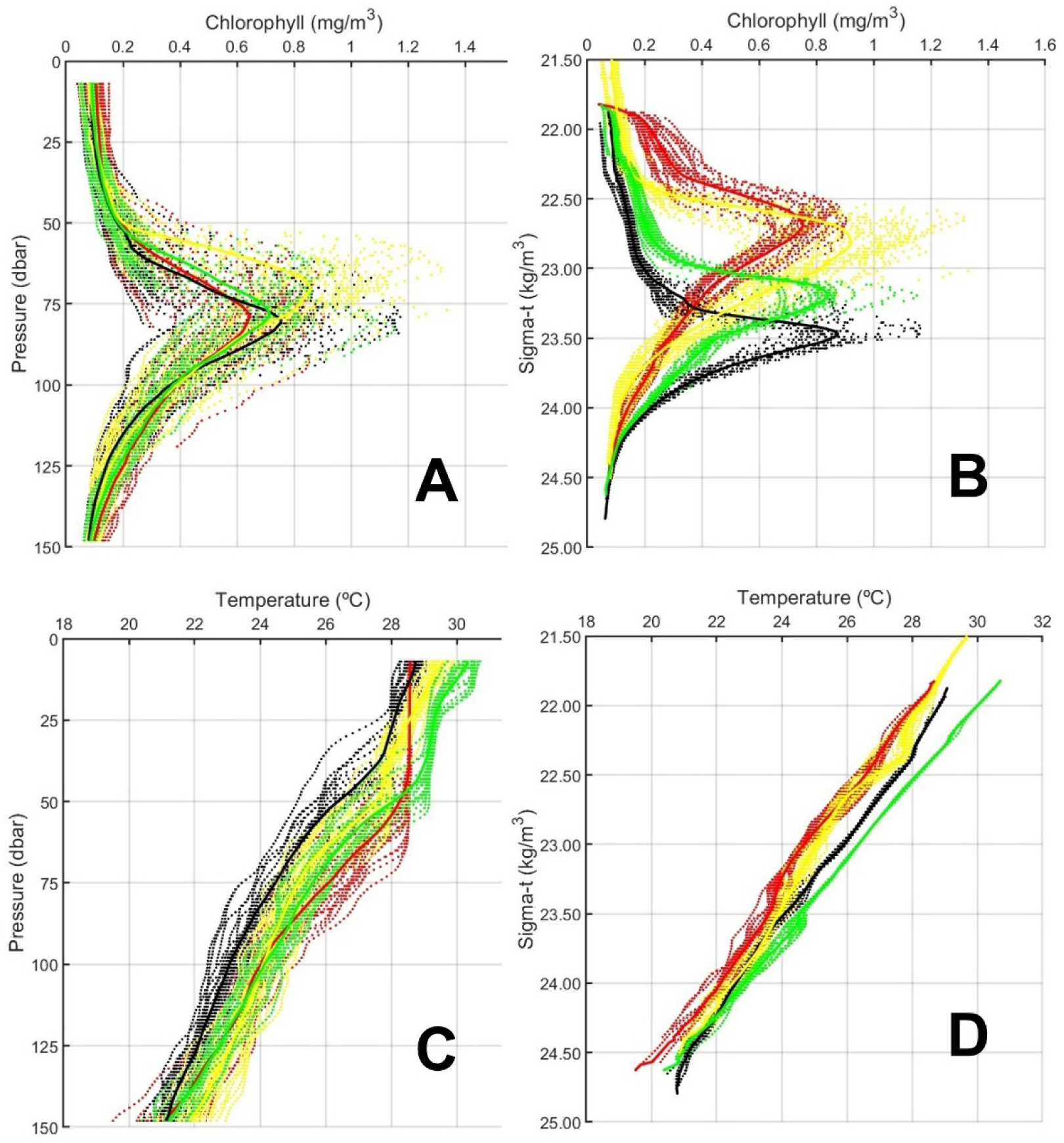
All chlorophyll and temperature vertical profiles as a function of pressure (A,C) and sigma-theta (B,D) for Cycle 1 (red dots), Cycle 2 (black dots), Cycle 3 (yellow dots) and Cycle 4 (green dots). Solid lines show the corresponding average vertical profiles.

**Figure 6.**
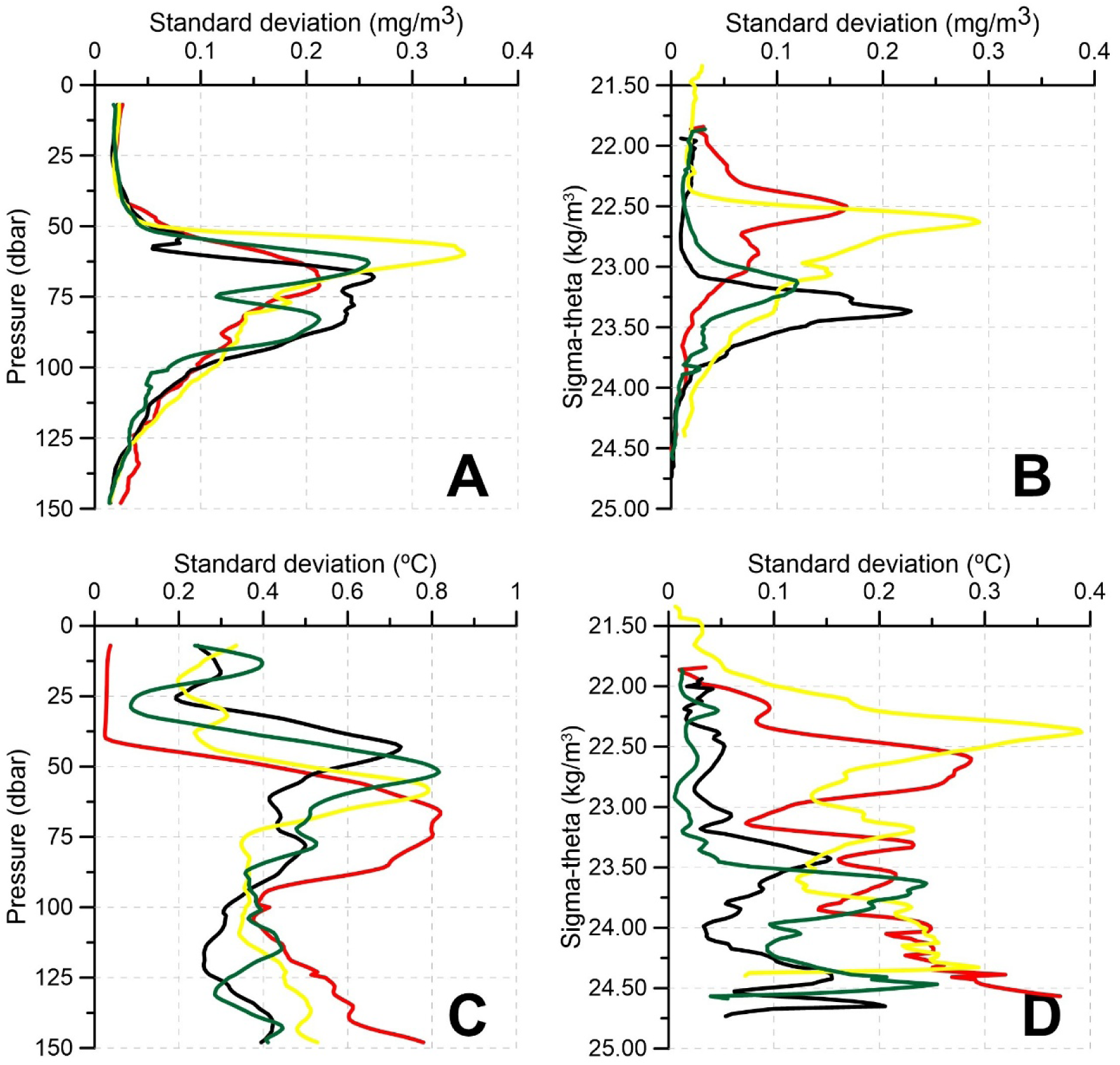
Standard deviations for chlorophyll (A,B) and temperature (C,D) as functions of pressure (A,C) and sigma-theta (B,D): Cycle 1 (red), Cycle 2 (black), Cycle 3 (yellow) and Cycle 4 (green). For each pressure level, all measurements from all cycle casts were used to calculate the standard deviation.

Two facts should be noticed in Figures 5 and 6. First, the dispersion of data points is much larger when chlorophyll and temperature data from the same cycle are compared at the same pressure level than for isopycnal surfaces. This is clearly reflected in the standard deviation reduction outlined in Figure 6. It is remarkable that when temperature is considered on sigma-theta levels, all the profiles from Cycles 1 and 4 are almost identical, while very small dispersion is observed for Cycles 2 and 3. The second fact is that the analyses on isopycnal surfaces, while showing lower differences within the same cycle, show larger differences when different cycles were compared (Figs. 5 A,B).

Shapiro-Wilk tests revealed that the distributions of the different variables analyzed did not follow normal distributions at most of the pressure or density levels. Therefore, Kruskal-Wallis tests were carried out at each vertical level (pressure or sigma-theta).

Figure 7 shows the H value (black solid line) for each pressure level for temperature (Fig. 7A), salinity (Fig. 7B), chlorophyll concentration (Fig. 7C) and dissolved oxygen concentration (Fig. 7D). Red solid lines are the H statistic calculated for each sigma- theta level and dashed lines give the critical values at the 0.05 significance level. These results show that the physical (temperature and salinity) and biochemical conditions (chlorophyll and oxygen concentrations) varied across the cycles.

**Figure 7.**
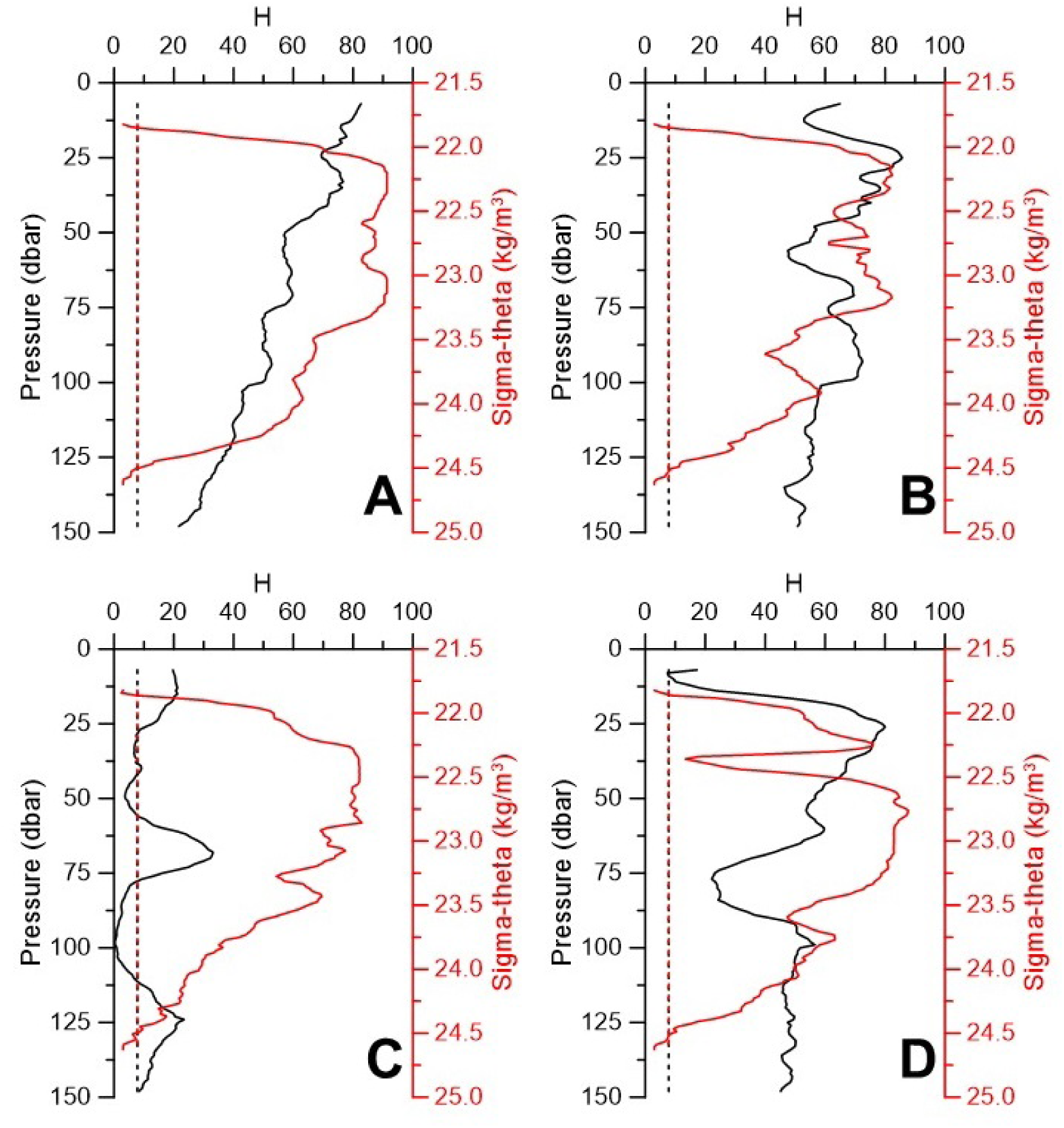
H statistics from Kruskal Wallis tests of the equality of the estimated mean values based on pressure (black lines) and sigma-theta (red lines) for the four cycles. A = temperature; B = salinity; C = chlorophyll; D = dissolved oxygen. Dashed lines show the critical values.

## 4. Discussion

In an Eulerian framework, sampling is conducted at fixed locations such as typically done with oceanographic stations distributed throughout a predefined study area.

Vertical profiles of physical and biogeochemical variables collected at multiple stations are then analyzed and compared to evaluate their spatial distributions, both horizontally and vertically. These analyses rely on the assumption of synoptic sampling. However, if the sampling period exceeds the characteristic time scales of the region’s oceanographic dynamics, this assumption may be invalidated, and the observed differences might reflect advective processes rather than true spatial variability.

Within this Eulerian approach, some oceanographic stations might be sampled repeatedly, for instance, in long-term time-series programs such as BATS (North Atlantic) and HOT-ALOHA (North Pacific) (Steinberg et al., 2001; Karl and Lukas, 1996). Although less common, stations may also be revisited during a single campaign. In both cases, there is a risk of misinterpreting advection-driven changes as temporal variations in physical or biogeochemical properties (Stukel et al., submitted).

To address the limitations of the Eulerian approach, a Lagrangian methodology can be applied. During the RR2201 cruise off the northwest coast of Australia, analyzed in this study, a drifter array was deployed to track a specific parcel of water. The water column was sampled at regular intervals over several days under the assumption that the measurements were being taken from the same moving water parcel.

Consider, for example, a water volume at 75-m depth. As this volume drifts with prevailing currents, its horizontal position changes over time. However, because the sampling location moves along with the water parcel, vertical profiles are consistently obtained from the same body of water. At 75 m, this means that the same phytoplankton and zooplankton communities inhabiting that layer are repeatedly sampled.

Consequently, any changes observed in these communities can be attributed to biological processes—such as net primary production, grazing, or bacterial degradation—rather than to advection. Similarly, variations in zooplankton or ichthyoplankton abundance could reflect in situ growth or mortality rates, given that the sampling point follows the drifting community. This approach assumes that water parcels predominantly drift horizontally, since horizontal currents are orders of magnitude stronger than vertical ones, as shown by the continuity equation (e.g., chapter 4 in Pond and Pickard, 1983, or any standard oceanography textbook).

The analysis of Cycle 1 casts 11 and 12 (Fig. 3A) revealed significant changes in chlorophyll and oxygen concentrations at fixed depths over a period of just three hours. For instance, chlorophyll concentration at 90 dbar increased from 0.4 to 0.8 mg m^-^³. If chlorophyll is used as a proxy for phytoplankton abundance, such a rapid increase might suggest an elevated net primary production rate. However, although extremely fast growth rates, with doubling times of less than a day, have been reported under optimal conditions (Sarmiento and Gruber, 2006), such rates are highly improbable in the oligotrophic waters of the eastern Indian Ocean (Davis et al., 2022; Kehinde et al., 2023). Conversely, chlorophyll concentration at 70 dbar decreased from 0.8 mg/m^3^ to 0.2 mg/m^3^, suggesting strong grazing pressure or rapid microbial degradation. Comparable discrepancies at fixed depths were also observed in dissolved oxygen, temperature, salinity, and density across most casts during the four experimental cycles.

Unlike biogeochemical variables, physical properties such as temperature, salinity, and density are conservative and, in the absence of mixing, should remain unchanged in a Lagrangian experiment. Mixing is more likely to occur in surface waters due to wind forcing. However, the uppermost 7 dbar of the water column were excluded from the present analysis to minimize this effect. Moreover, wind speeds during the four cycles remained below 6 m/s and exhibited a decreasing trend, reaching approximately 2 m/s by the middle of Cycle 3 (Fig. 8A). The wind predominantly blew from the southeast, with occasional interruptions, consistent with the year-round prevalence of southeasterly trade winds south of 10°S (Phillips et al., 2021). A stick diagram of the wind (Fig. 8B) shows that wind vectors were directed toward the northwest. This observation is supported by monthly averaged wind data from the COPERNICUS dataset (Supplementary Fig. S3).

**Figure 8.**
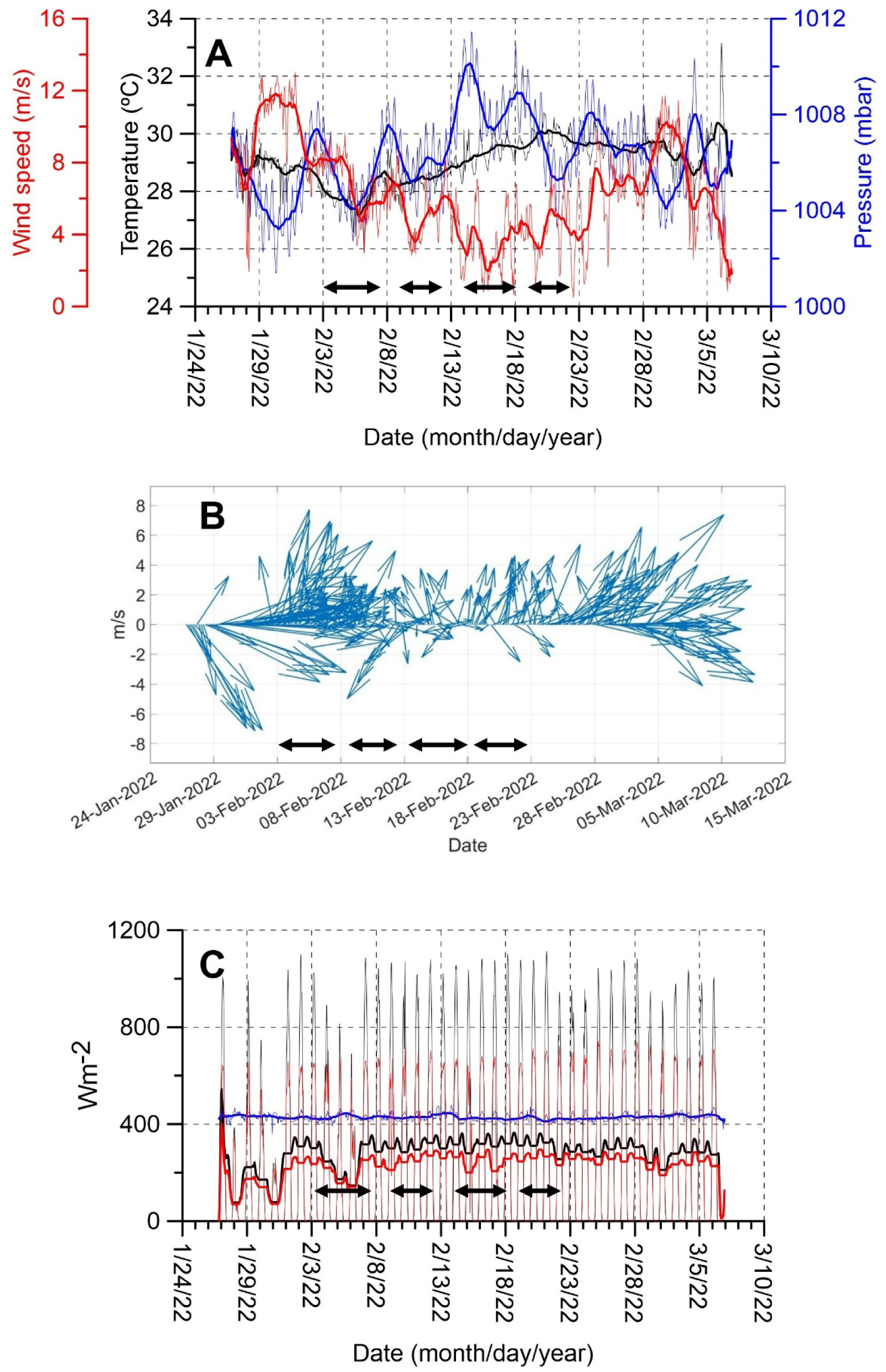
Meteorological and sea surface data recorded by the onboard station of R/V Roger Revelle during cruise RR2201. Panel 8A shows wind speed (red line), atmospheric pressure (blue line), and air temperature (black line). Figure 8B shows wind vectors. Figure 8C displays incident shortwave radiation (black line), photosynthetically active radiation (PAR, blue line), and emitted longwave radiation (red line).

The most significant changes in both physical and biogeochemical variables were observed below the surface mixed layer, particularly between 50 and 150 dbar (Fig. 3D). This further suggests that wind-driven mixing was an unlikely explanation for the variability observed between consecutive casts, but it raises the question of whether vertical shear in the water column may have caused the deeper layers to follow a different trajectory than that followed by the drogue within the surface mixed-layer.

However, an examination of the temporal evolution of vertical density profiles (Figs. 4 A,F,K and P) reveals continuous vertical displacements of isopycnals, indicative of internal tide activity. In some instances, isopycnals shifted by more than 20 m between consecutive casts (Fig. 4). Such vertical movements imply the displacement of water parcels, along with their resident phytoplankton and zooplankton communities and associated biogeochemical properties, up and down in the water column. Figure 4 shows that these vertical excursions were more evident below 50 dbar. Consequently, the conservative character of density is an acceptable assumption that allows us to calculate vertical displacements of water parcels and to decompose the between-cast depth variability into contributions from vertical movements (heaving) and changes on isopycnals.

This decomposition demonstrated that the largest contribution to variability observed on depth levels was from vertical movements of water parcels (Fig. 3). Consider a water parcel with a given chlorophyll and oxygen concentration that occupy a certain depth level in a cast. If this water parcel experiences an upward displacement of 20 m, during the following cast, we would find those same values of chlorophyll and oxygen concentrations 20 m higher up, unless some biochemical processes alter them. Therefore, the changes in chlorophyll and oxygen concentrations corresponding to the same density value between two consecutive casts could be attributed to processes such as net primary production, bacterial degradation or simply photo-acclimation. These changes on isopycnals in the decomposition outlined in Subsection 2.3 is an adaptation of Bindoff and McDougal (1994). However, in some cases, the magnitude of these changes may be too large to be explained solely by biochemical processes. For instance, a sharp increase in chlorophyll and oxygen concentrations over just a few hours would be highly unlikely in oligotrophic waters.

As previously noted, the results presented in Figure 3 show that changes along isopycnals are significantly smaller than those observed on isobaric levels, with the latter being primarily driven by vertical displacements (heaving). Moreover, when using sigma-theta as the vertical coordinate to analyze the temporal evolution of chlorophyll and oxygen concentrations (Fig. 4 and S1), as well as temperature and salinity (Figure S2, supplementary material), the isolines become noticeably flatter, often nearly horizontal. This indicates that when following the evolution of these properties within water parcels and accounting for their vertical displacements, these variables exhibit minimal short-term variation over the duration of each experimental cycle.

The decomposition applied in this study confirms that the four cycles conducted during the RR2201 cruise were indeed Lagrangian in nature, with the same water parcel effectively tracked and sampled throughout. Additionally, the analysis reveals that biogeochemical changes were relatively small and that stable conditions prevailed within each cycle. The lack of substantial changes in biogeochemical properties along isopycnals, combined with the fact that most of the observed variability could be attributed to vertical movement of water masses, strongly supports the Lagrangian character of the sampling.

However, the reverse is not necessarily true. Significant changes along isopycnals may still occur, as biogeochemical variables are not conservative and can be influenced by various processes within a single water parcel. In such cases, other factors—such as the time scales over which these changes occur and their plausibility based on existing oceanographic knowledge of the study region—must be taken into account. In other words, confirming the Lagrangian nature of an experiment is generally more straightforward than disproving it.

In addition to confirming that the same water parcel was sampled throughout each cycle, Figure 5D and S2 demonstrate that identical temperature profiles were obtained across all casts within the same cycle, indicating stable physical conditions throughout the duration of each cycle. This stability was also observed for chlorophyll and oxygen concentrations (Fig. 4). Such consistency likely reflects the absence of strong mixing events during the cycles (Fig. 8), the short duration of each cycle (3 to 4 days), and the prevailing oligotrophic conditions.

We note that the small changes observed in chlorophyll and oxygen concentrations along isopycnal surfaces do not seem to match day-night cycles (see midnight and noon times along the cycles in Figure S1 in supplementary material). Nor do they seem to be the result of changes in net primary production, in which case, we might expect a positive correlation between net increase of chlorophyll and an increased dissolved oxygen concentration. This expectation might, however, oversimplify the more complex dynamics of living cells and communities as they respond to changes in their physical and nutritional environments. Cellular chlorophyll content, for example, is actively regulated by living phytoplankton with respect to the light environment and should be upregulated when internal waves drive the cells deeper in the water column and downregulated when the cells are at shallower depths. Similarly, variations in oxygen concentration do not reflect the instantaneous balance of photosynthesis and respiration rates, but the cyclical behavior of substrate production and microbial respiratory responses to substrate availability which might have its own rhythm in response to internal wave perturbations. Figure 3H shows these changes for Cycle 1 calculated along isopycnals (therefore likely to biological interactions) and the same information can be seen for the four cycles in Figure S4. This is an interesting question that requires further investigation, but is out of the scope of the present work.

The stability of physical and biochemical conditions within each cycle had important implications for the experimental design. It allowed us to summarize the environmental conditions of each cycle by computing mean profiles of temperature, salinity, chlorophyll, and oxygen concentrations. For each depth level and variable, we assumed that the values obtained from different casts within the same cycle originated from a single statistical distribution, characterized by its mean and standard deviation (Figs. 5 and 6).

The next step was to evaluate, for each variable and at each pressure or density level, whether the four distributions (corresponding to the four cycles) followed a normal distribution, and whether differences among them were statistically significant. The Shapiro-Wilk and Kruskal-Wallis tests rejected normality and confirmed significant differences between cycles. Although relating these differences to tuna larval growth rates or their environmental suitability is beyond the scope of the present work, such aspects will be addressed in Laiz et al. (this volume). It is important to note, however, that these analyses would not have been possible without first establishing the Lagrangian character of the sampling and the environmental stability within each cycle.

Naturally, had our results indicated that the casts within a single cycle did not originate from the same water column, it would have been meaningless to average the profiles to represent environmental conditions for that cycle. Likewise, if significant changes in physical or biogeochemical properties had been observed along the parcel’s trajectory due to strong mixing, high net primary production, or intense phytoplankton grazing, it would not have been appropriate to characterize the cycle using averaged profiles, as temporal evolution would need to be accounted for.

## 5. Conclusions

In summary, when repeated sampling is conducted over time, observed changes on depth (isobaric) levels can be decomposed into two contributions: vertical displacements of water masses (heaving), and changes along isopycnal surfaces. The latter reflects intrinsic changes within a water parcel and, in the case of biogeochemical variables, may be linked to biological processes affecting phytoplankton and zooplankton communities. This decomposition method is well suited for Lagrangian experiments such as the one conducted during the BLOOFINZ cruise to study the trophic ecology of southern bluefin tuna (SBT) larvae in the only known spawning ground off northwest Australia. In this case, the methodology helps to confirm the Lagrangian character of the experiments and reveals significant environmental differences among cycles. This information is essential to better understand larval habitat quality in these oligotrophic habitats and how the energy is transferred from the base of the food web up to the larva, with direct implications in larval growth and survival, crucial information for ecosystem-based fisheries management. This approach could also be applied to other studies involving repeated sampling at oceanographic stations, such as yo-yo stations.

## Supporting information

Position of casts

## Declaration of Competing Interest

The authors declare no personal or financial conflicts of interest that could have influenced the results or conclusions of this study.

## Author Statement

## Some material for Acknowledgements

We sincerely thank the crew of the R/V Roger Revelle for their assistance during the RR2201 survey. Cruise RR2201 was supported by U.S. National Science Foundation grants [OCE-1851558](https://www.bco-dmo.org/award/819487) (M.R.L.), -1851347 and -2404504 (S.A.K. and M.R.S) and is a contribution to the Second International Indian Ocean Expedition (IIOE-2 endorsed project EP046). This study was partialy funded by the INDITUN project PID2021/122862NB/100 UE-FEDER (R.L.-C.) supported by the Ministry of Science, Innovation and Universities (MICINN) of the Spanish government. Seawater sampling was done under Australian Government permit AU-COM2021-520 and Australian Marine Parks permit PA2021-00062-2 issued by the Director of National Parks, Australia. Views expressed in this publication do not necessarily represent those of the Director of National Parks or the Australian Government.

## Notes

### Competing Interest Statement

The authors have declared no competing interest.

## References

Bindoff, N.L., McDougall, T.J., 1994. Diagnosing climate change and ocean ventilation using hydrographic data. Journal of Physical Oceanography, 24, 1137–1152. 10.1175/1520-0485(1994)024<1137:DCCAOV>2.0.CO;2.

Bornatowski, H., Angelini, R., Coll, M., Barreto, R. R. P., Amorim, A. F., 2017. Ecological role and historical trends of large pelagic predators in a subtropical marine ecosystem of the South Atlantic. Reviews in Fish Biology and Fisheries, 28, 241–259. 10.1007/s11160-017-9492-z.

Estes, J.A., Terborgh, J., Brashares, J.S., Power, M.E., Berger, J., Bond, W. J., Carpenter, S.R., Essington, T.E., Holt, R.D., Jackson, J.B.C., Marquis, R.J., Oksanen, L., Oksanen, T., Paine, R.T., Pikitch, E.K., Ripple, W.J., Sandin, S.A., Scheffer, M., Schoener, T.W., Shurin, J.B., Sinclair, A.R.E., Soulé, M.E., Virtanen, R., Wardle, D.A., 2011. Trophic Downgrading of Planet Earth. Science, 333(6040):301–306. https://www.science.org/doi/10.1126/science.1205106.

Davies, C. H., Beckley, L. E., Richardson, A. J., 2022. Copepods and mixotrophic Rhizaria dominate zooplankton abundances in the oligotrophic Indian Ocean. Deep-Sea Res. II, 202, 105136. 10.1016/j.dsr2.2022.105136.

Davis, T. L. O., Jenkins, G. P., Young, J. W., 1990. Diel patterns of vertical distribution in larvae of southern bluefin *Thunnus maccoyii*, and other tuna in the East Indian Ocean. Mar. Ecol. Prog. Ser., 59, 63–74.

FAO. 2024. The State of World Fisheries and Aquaculture 2024. Blue Transformation in action. Rome. 10.4060/cd0683en.

Gerard, T., Lamkin, J., Kelly, T. B., Knapp, A. N., Laiz-Carrión, R., Malca, E., Selph, K. E., Shiroza, A., Shropshire, T. A., Stukel, M. R., Swalethorp, R., Yingling, N., Landry, M. R., 2022. Bluefin Larvae in Oligotrophic Ocean Foodwebs, investigations of nutrients to zooplankton: overview of the BLOOFINZ-Gulf of Mexico program. Journal of Plankton Research, 44, Issue 5, September/October 2022, Pages 600-617, 10.1093/plankt/fbac038.

Holm-Hansen, O., Lorenzen, C.J., Holmes, R.W., Strickland, J.D., 1965. Fluorometric determination of chlorophyll. J. Mar. Sci. 30 (1), 3–15, 10.1093/icesjms/30.1.3.

ISSF. 2022. Status of the world fisheries for tuna. Mar. 2022. ISSF Technical Report 2022-04. International Seafood Sustainability Foundation, Washington, D.C., USA.

Jeffrey, S. W., Humphrey, G. F., 1975. New spectrophotometric equations for determining chlorophylls a, b, c1 and c2 in higher plants, algae and natural phytoplankton. Biochemie und Physiologie der Pflanzen, *167* (2), 191–194. 10.1016/S0015-3796(17)30778-3.

Jenkins, G. P., Davis, T. L. O., 1990. Age, growth rate, and growth trajectory determined from otolith microstructure of southern bluefin tuna *Thunnus maccoyii* larvae. Mar. Ecol. Prog. Ser., 63, 93–104.

Karl, D.M., Lukas, R., 1996. The Hawaii Ocean Time-series (HOT) program: Background, rationale and field implementation. Deep Sea Res. Part II Top. Stud. Oceanogr., 43, 129–156.

Kehinde, O., Bourassa, M., Kranz, S., Landry, M. R., Kelly, T., Stukel, M. R., 2023. Lateral advection of particulate organic matter in the eastern Indian Ocean. Journal of Geophysical Research: Oceans, 128, e2023JC019723. 10.1029/2023JC019723

Kelly, T.B., Knapp, A.N., Landry, M.R., Selph, K. E., Shropshire, T. A., Thomas, R. K., Stukel, M. R., 2021. Lateral advection supports nitrogen export in the oligotrophic open-ocean Gulf of Mexico. Nat Commun 12, 3325 10.1038/s41467-021-23678-9

Kim, Y.-S., Delgado, D. Y., Cano, I. A., Sawada, Y., 2015. Effect of temperature and salinity on hatching and larval survival of yellowfin tuna *Thunnus albacares*. Fisheries Science 81(5). *Fish Sci.* 10.1007/s12562-015-0901-8.

Laiz-Carrión, R., Gerard, T., Suca, J. J., Malca, E., Uriarte, A., Quintanilla, J. M., Privoznik, S., Llopiz, J. K., Lamkin, J., García, A., 2019. Stable isotope analysis indicates resource partitioning and trophic niche overlap in larvae of four tuna species in the Gulf of Mexico. Mar. Ecol. Prog. Ser, 619: 53–68, 10.3354/meps12958.

Nieblas, A.-E., Demarcq, H., Drushka, K., Sloyan, B., Bonhommeau, S., 2014. Front variability and surface ocean features of the presumed southern bluefin tuna spawning grounds in the tropical southeast Indian Ocean. Deep-Sea Research II, 107, 64–76. 10.1016/j.dsr2.2013.11.007.

Phillips, H. E., Tandon, A., Furue, R., Hood, R., Ummenhofer, C. C., Benthuysen, J. A., Menezes, V., Hu, S., Webber, B., Sanchez-Franks, A., Cherian, D., Shroyer, E., Feng, M., Wijesekera, H., Chatterjee, A., Yu, L., Hermes, J., Murtugudde, R., Tozuka, T., Su, D., Singh, A., Centurioni, L., Prakash, S., Wiggert, J., 2021. Progress in understanding of Indian Ocean circulation, variability, air–sea exchange, and impacts on biogeochemistry, Ocean Sci., 17, 1677–1751. 10.5194/os-17-1677-2021.

Pond, S., Pickard, G. L., 1983. Introductory Dynamical Oceanography. Second edition. Pergamon Press*. Oxford*.

Quintanilla, J.M., Borrego-Santos, R., Malca, E., Swalethorp, R., Landry, M.R., Gerard, T., Lamkin, J., García, A., Laiz-Carrión, R., 2024. Maternal Effects and Trophodynamics Drive Interannual Larval Growth Variability of Atlantic Bluefin Tuna (*Thunnus thynnus)* from the Gulf of Mexico. Animals, 14, 1319. 10.3390/ani14091319.

Quintanilla, J. M., Malca, E., Lamkin, J., García, A., Laiz-Carrión, R., 2023. Evidence of isotopic maternal transmission influence on bluefin tuna (*Thunnus thynnus*) larval growth. Marine Environmental Research, 190, 106112. 10.1016/j.marenvres.2023.106112.

Sarmiento, J. L., Gruber, N., 2006. Ocean Biogeochemical Dynamics. Princeton University Press. 10.2307/j.ctt3fgxqx.

Satoh, K., Tanaka, Y., Iwahashi, M., 2008. Variations in the instantaneous mortality rate between larval patches of Pacific bluefin tuna *Thunnus orientalis* in the northwestern *Pacific Ocean*. Fisheries Research 89, 248–256. 10.1016/j.fishres.2007.09.003.

Steinberg, D.K., Carlson, C.A., Bates, N.R., Johnson, R.J., Michaels, A.F., Knap, A.H., 2001. Overview of the US JGOFS Bermuda Atlantic Time-series Study (BATS): A decade-scale look at ocean biology and biogeochemistry. Deep Sea Res. Part II Top. Stud. Oceanogr., 48, 1405–1447.

Strickland, J.D.H., Parsons, T., 1972. A practical handbook of seawater analysis. B. Fish. Res. Board Can. 167, 293 pp. 10.1002/iroh.19700550118.

Stukel, M. R., Allen, A. E., Barbeau, K. A., Chabert, P., Dovel, S., Gangrade, S., Kranz, S. A., Lampe, R. H., Landry, M. R., Marrec, P., Messié, M., Miller, A. J., Wilkinson, G., Ohman, M. D. Submitted to BioScience.

Wexler, J. B., Margulies, D., Scholey, V. P., 2011. Temperature and dissolved oxygen requirements for survival of yellowfin tuna, Thunnus albacares, larvae. Journal of Experimental Marine Biology and Ecology. 404, 63–72, 10.1016/j.jembe.2011.05.002.

