## Supplementary material for "Analysis of environmental data during a lagrangian experiment: The influence of vertical movements": Position of casts

Table S1. Cast number, longitude, latitude, year, month, day, hour and minute for each of the CTD casts corresponding to the first cycle.

| **CYCLE 1** | | | | | | | |
| --- | --- | --- | --- | --- | --- | --- | --- |
| **Cast** | **Longitude** | **Latitude** | **Year** | **Month** | **Day** | **Hour** | **Minute** |
| 11 | 114,55 | -15,35 | 2022 | 2 | 3 | 19 | 26 |
| 12 | 114,50 | -15,33 | 2022 | 2 | 3 | 22 | 30 |
| 13 | 114,50 | -15,33 | 2022 | 2 | 3 | 23 | 24 |
| 14 | 114,51 | -15,35 | 2022 | 2 | 4 | 4 | 32 |
| 15 | 114,51 | -15,35 | 2022 | 2 | 4 | 5 | 37 |
| 16 | 114,43 | -15,37 | 2022 | 2 | 4 | 12 | 33 |
| 17 | 114,40 | -15,37 | 2022 | 2 | 4 | 14 | 45 |
| 18 | 114,37 | -15,37 | 2022 | 2 | 4 | 18 | 27 |
| 19 | 114,36 | -15,37 | 2022 | 2 | 4 | 21 | 46 |
| 20 | 114,36 | -15,37 | 2022 | 2 | 4 | 23 | 9 |
| 21 | 114,34 | -15,36 | 2022 | 2 | 5 | 4 | 26 |
| 22 | 114,34 | -15,36 | 2022 | 2 | 5 | 5 | 19 |
| 23 | 114,27 | -15,37 | 2022 | 2 | 5 | 12 | 32 |
| 24 | 114,27 | -15,37 | 2022 | 2 | 5 | 14 | 7 |
| 25 | 114,23 | -15,40 | 2022 | 2 | 5 | 18 | 5 |
| 26 | 114,22 | -15,40 | 2022 | 2 | 5 | 21 | 43 |
| 27 | 114,22 | -15,40 | 2022 | 2 | 5 | 22 | 42 |
| 28 | 114,20 | -15,43 | 2022 | 2 | 6 | 4 | 50 |
| 29 | 114,20 | -15,44 | 2022 | 2 | 6 | 5 | 56 |
| 30 | 114,19 | -15,45 | 2022 | 2 | 6 | 7 | 29 |
| 31 | 114,18 | -15,48 | 2022 | 2 | 6 | 12 | 23 |
| 32 | 114,18 | -15,48 | 2022 | 2 | 6 | 13 | 50 |
| 33 | 114,19 | -15,47 | 2022 | 2 | 6 | 18 | 28 |
| 34 | 114,18 | -15,47 | 2022 | 2 | 6 | 21 | 28 |
| 35 | 114,18 | -15,47 | 2022 | 2 | 6 | 22 | 34 |
| 36 | 114,16 | -15,50 | 2022 | 2 | 7 | 4 | 49 |
| 38 | 114,14 | -15,60 | 2022 | 2 | 7 | 19 | 9 |

Table S2. Cast number, longitude, latitude, year, month, day, hour and minute for each of the CTD casts corresponding to the second cycle.

| **CYCLE 2** | | | | | | | |
| --- | --- | --- | --- | --- | --- | --- | --- |
| **Cast** | **Length** | **Latitude** | **Year** | **Month** | **Day** | **Hour** | **Minute** |
| 47 | 115,69 | -16,73 | 2022 | 2 | 9 | 14 | 20 |
| 48 | 115,71 | -16,75 | 2022 | 2 | 9 | 19 | 4 |
| 49 | 115,75 | -16,74 | 2022 | 2 | 9 | 21 | 49 |
| 50 | 115,75 | -16,74 | 2022 | 2 | 9 | 22 | 33 |
| 51 | 115,76 | -16,74 | 2022 | 2 | 10 | 5 | 21 |
| 52 | 115,76 | -16,74 | 2022 | 2 | 10 | 6 | 17 |
| 54 | 115,83 | -16,82 | 2022 | 2 | 10 | 18 | 32 |
| 55 | 115,85 | -16,83 | 2022 | 2 | 10 | 21 | 45 |
| 56 | 115,85 | -16,83 | 2022 | 2 | 10 | 23 | 6 |
| 57 | 115,91 | -16,83 | 2022 | 2 | 11 | 4 | 31 |
| 58 | 115,91 | -16,83 | 2022 | 2 | 11 | 6 | 9 |
| 59 | 115,93 | -16,84 | 2022 | 2 | 11 | 12 | 12 |
| 60 | 115,94 | -16,86 | 2022 | 2 | 11 | 14 | 6 |
| 61 | 115,94 | -16,88 | 2022 | 2 | 11 | 18 | 4 |
| 62 | 115,95 | -16,92 | 2022 | 2 | 11 | 21 | 41 |
| 63 | 115,95 | -16,92 | 2022 | 2 | 11 | 22 | 49 |
| 64 | 115,99 | -16,95 | 2022 | 2 | 12 | 5 | 28 |
| 65 | 116,01 | -16,97 | 2022 | 2 | 12 | 7 | 38 |
| 66 | 116,05 | -16,99 | 2022 | 2 | 12 | 12 | 17 |
| 67 | 116,05 | -17,01 | 2022 | 2 | 12 | 13 | 44 |
| 68 | 116,07 | -17,00 | 2022 | 2 | 12 | 18 | 5 |
| 69 | 116,08 | -17,01 | 2022 | 2 | 12 | 21 | 36 |
| 70 | 116,08 | -17,01 | 2022 | 2 | 12 | 22 | 39 |

Table S3. Cast number, longitude, latitude, year, month, day, hour and minute for each of the CTD casts corresponding to the third cycle.

| **CYCLE 3** | | | | | | | |
| --- | --- | --- | --- | --- | --- | --- | --- |
| **Cast** | **Length** | **Latitude** | **Year** | **Month** | **Day** | **Hour** | **Minute** |
| 75 | 115,59 | -16,10 | 2022 | 2 | 14 | 7 | 11 |
| 76 | 115,61 | -16,00 | 2022 | 2 | 14 | 10 | 4 |
| 77 | 115,52 | -16,10 | 2022 | 2 | 14 | 13 | 5 |
| 78 | 115,65 | -16,17 | 2022 | 2 | 14 | 16 | 13 |
| 79 | 115,78 | -16,06 | 2022 | 2 | 14 | 19 | 15 |
| 80 | 115,69 | -15,87 | 2022 | 2 | 14 | 22 | 19 |
| 81 | 115,81 | -16,02 | 2022 | 2 | 15 | 2 | 39 |
| 82 | 115,91 | -16,02 | 2022 | 2 | 15 | 5 | 43 |
| 83 | 115,83 | -15,98 | 2022 | 2 | 15 | 10 | 8 |
| 84 | 115,83 | -15,98 | 2022 | 2 | 15 | 11 | 54 |
| 85 | 115,83 | -15,98 | 2022 | 2 | 15 | 14 | 22 |
| 86 | 115,83 | -15,94 | 2022 | 2 | 15 | 18 | 29 |
| 87 | 115,82 | -15,93 | 2022 | 2 | 15 | 21 | 8 |
| 88 | 115,82 | -15,94 | 2022 | 2 | 15 | 22 | 19 |
| 89 | 115,83 | -15,92 | 2022 | 2 | 16 | 4 | 35 |
| 90 | 115,83 | -15,92 | 2022 | 2 | 16 | 5 | 41 |
| 91 | 115,83 | -15,90 | 2022 | 2 | 16 | 12 | 5 |
| 92 | 115,84 | -15,90 | 2022 | 2 | 16 | 13 | 51 |
| 93 | 115,84 | -15,88 | 2022 | 2 | 16 | 18 | 10 |
| 94 | 115,84 | -15,86 | 2022 | 2 | 16 | 21 | 9 |
| 95 | 115,83 | -15,83 | 2022 | 2 | 17 | 4 | 43 |
| 96 | 115,83 | -15,83 | 2022 | 2 | 17 | 5 | 56 |
| 97 | 115,82 | -15,81 | 2022 | 2 | 17 | 10 | 39 |
| 98 | 115,82 | -15,78 | 2022 | 2 | 17 | 14 | 5 |
| 99 | 115,80 | -15,76 | 2022 | 2 | 17 | 18 | 5 |
| 100 | 115,80 | -15,75 | 2022 | 2 | 17 | 21 | 8 |
| 101 | 115,80 | -15,75 | 2022 | 2 | 17 | 22 | 20 |
| 102 | 115,77 | -15,70 | 2022 | 2 | 18 | 4 | 45 |
| 103 | 115,77 | -15,70 | 2022 | 2 | 18 | 6 | 18 |
| 104 | 115,77 | -15,69 | 2022 | 2 | 18 | 8 | 27 |

Table S4. Cast number, longitude, latitude, year, month, day, hour and minute for each of the CTD casts corresponding to the fourth cycle.

| **CYCLE 4** | | | | | | | |
| --- | --- | --- | --- | --- | --- | --- | --- |
| **Cast** | **Length** | **Latitude** | **Year** | **Month** | **Day** | **Hour** | **Minute** |
| 110 | 117,99 | -16,00 | 2022 | 2 | 19 | 22 | 39 |
| 111 | 118,14 | -15,89 | 2022 | 2 | 20 | 10 | 22 |
| 112 | 118,14 | -15,89 | 2022 | 2 | 20 | 11 | 49 |
| 113 | 118,14 | -15,89 | 2022 | 2 | 20 | 13 | 52 |
| 114 | 118,14 | -15,89 | 2022 | 2 | 20 | 18 | 21 |
| 115 | 118,15 | -15,88 | 2022 | 2 | 20 | 20 | 46 |
| 116 | 118,14 | -15,88 | 2022 | 2 | 20 | 21 | 34 |
| 117 | 118,13 | -15,89 | 2022 | 2 | 21 | 4 | 31 |
| 118 | 118,13 | -15,89 | 2022 | 2 | 21 | 5 | 42 |
| 119 | 118,09 | -15,90 | 2022 | 2 | 21 | 11 | 44 |
| 120 | 118,10 | -15,90 | 2022 | 2 | 21 | 13 | 45 |
| 121 | 118,09 | -15,91 | 2022 | 2 | 21 | 18 | 10 |
| 122 | 118,09 | -15,93 | 2022 | 2 | 21 | 21 | 28 |
| 123 | 118,09 | -15,93 | 2022 | 2 | 21 | 22 | 27 |
| 124 | 118,11 | -15,95 | 2022 | 2 | 22 | 4 | 38 |
| 125 | 118,11 | -15,95 | 2022 | 2 | 22 | 5 | 49 |
| 126 | 118,11 | -15,95 | 2022 | 2 | 22 | 11 | 2 |
| 127 | 118,12 | -15,96 | 2022 | 2 | 22 | 14 | 22 |
| 128 | 118,12 | -15,94 | 2022 | 2 | 22 | 18 | 8 |

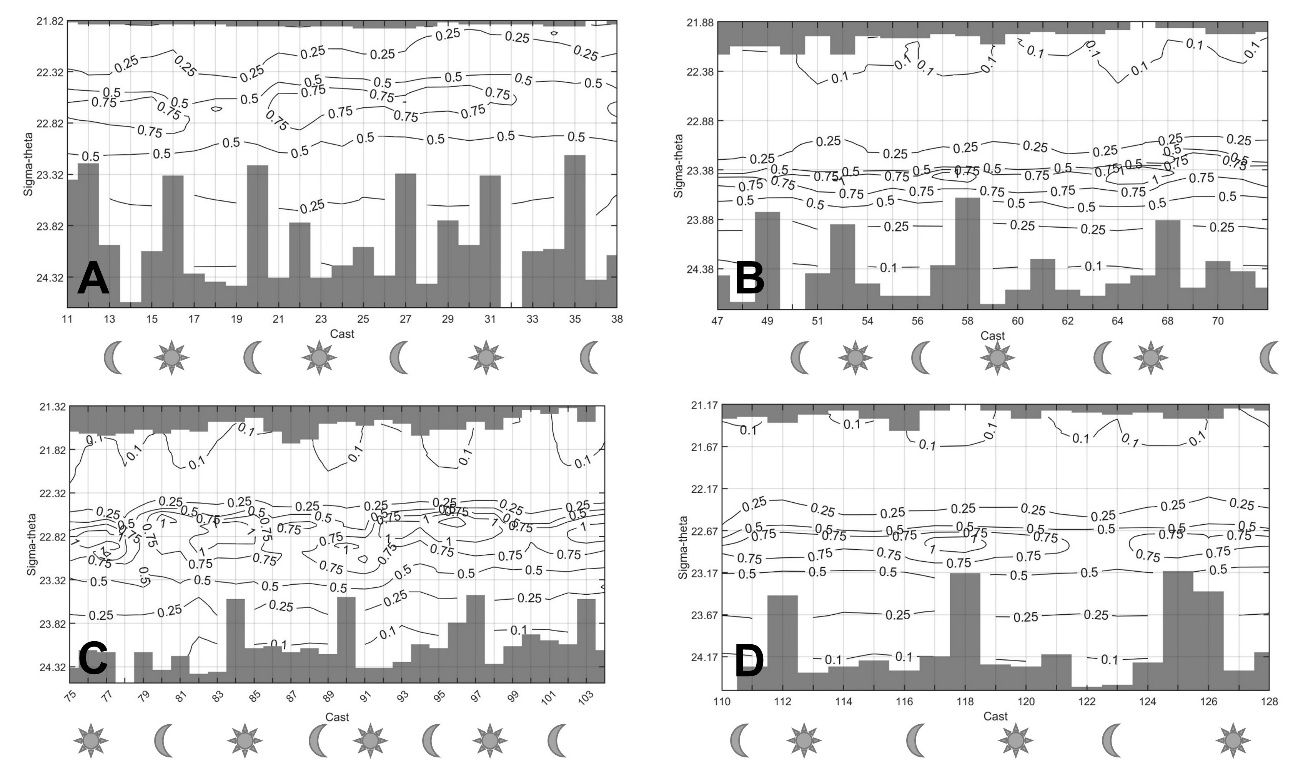

Figure S1. Hovmöller diagrams for the temporal evolution of the vertical dependence of chlorophyll concentration for cycles 1 (A), 2(B), 3(C) and 4(D). Sigma-theta has been used as vertical coordinate.

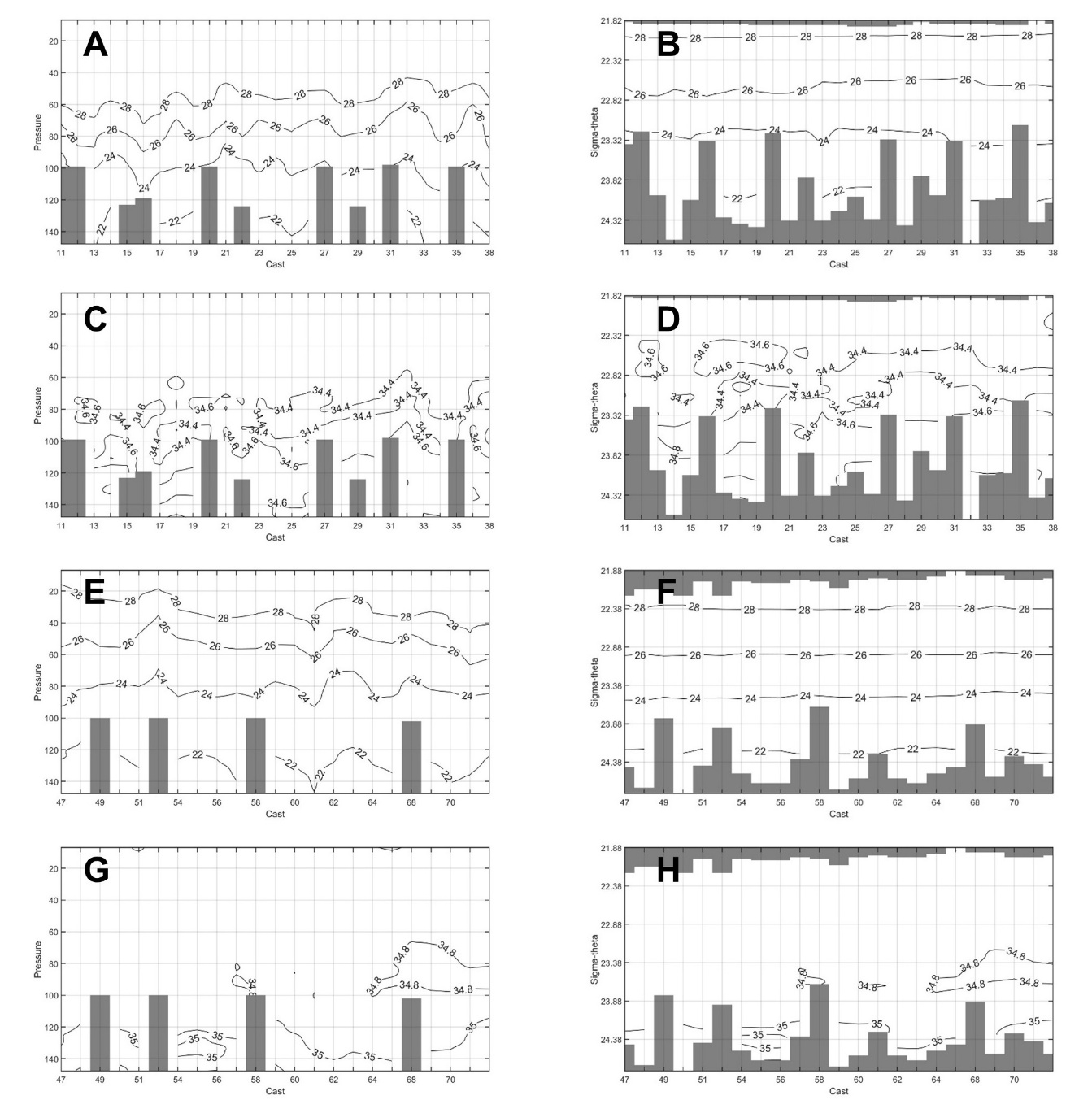

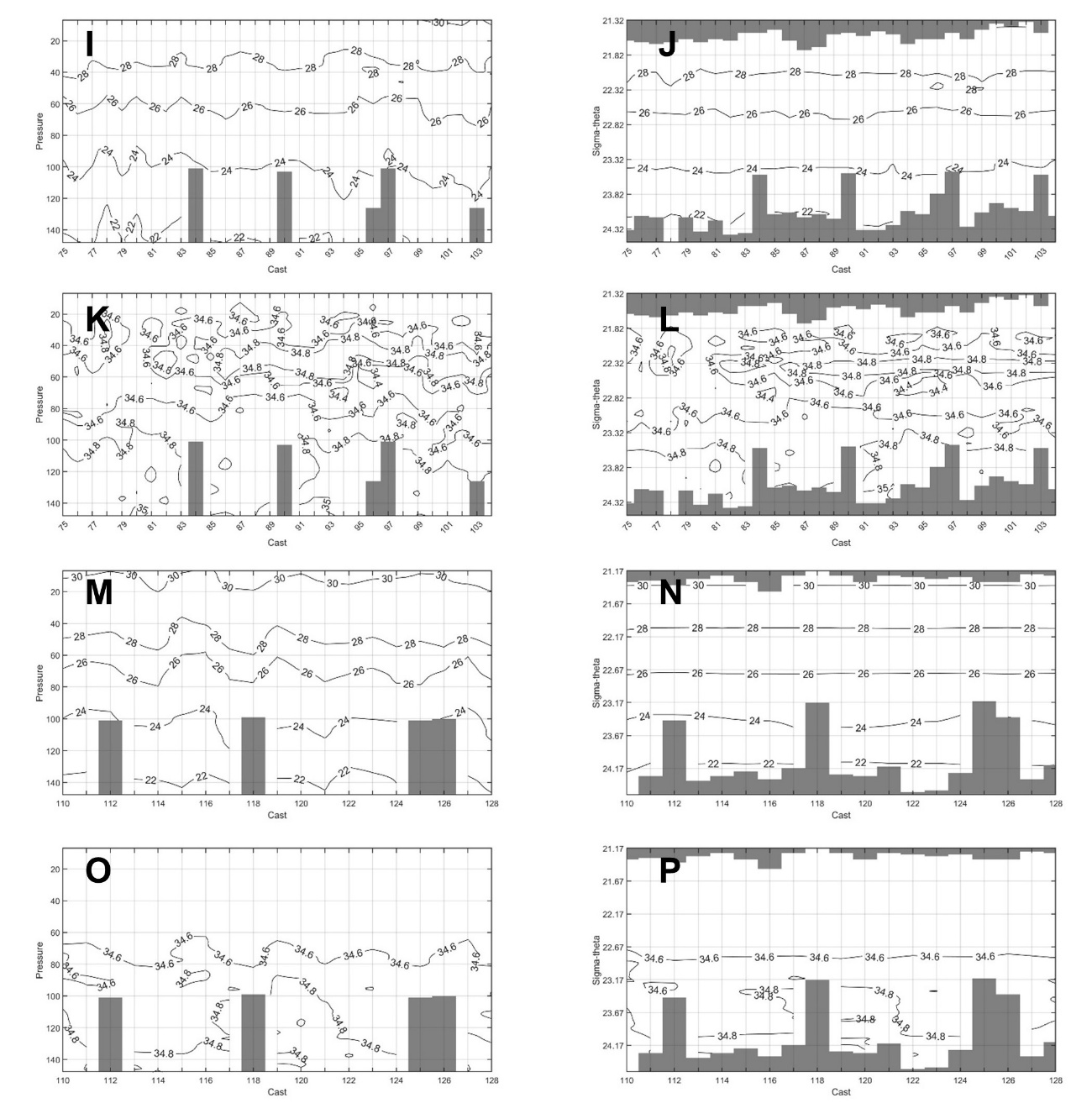

Figure S1. Figures S1A and S1B show the time evolution of the temperature for the first cycle. Figure S1A uses pressure as the vertical coordinate whereas sigma-theta is the vertical coordinate in Figure S1B. Figures S1C and S1D show the time evolution of salinity for the first cycle, using pressure (Fig. S1C) and sigma-theta (Fig. S1D) as vertical coordinates. Figures S1E and F show similar results for the evolution of temperature during the second cycle and Figures S1G and S1H correspond to the salinity of the second cycle. Notice that, in all the cases, the cast numbers were used as horizontal coordinates. Since consecutive casts were not always separated by the same time interval, the x-axes do not have a regular time step. Figures S1I to S1L show the time evolution of temperature and salinity for the third cycle and figures S1M to S1P correspond to the fourth cycle.

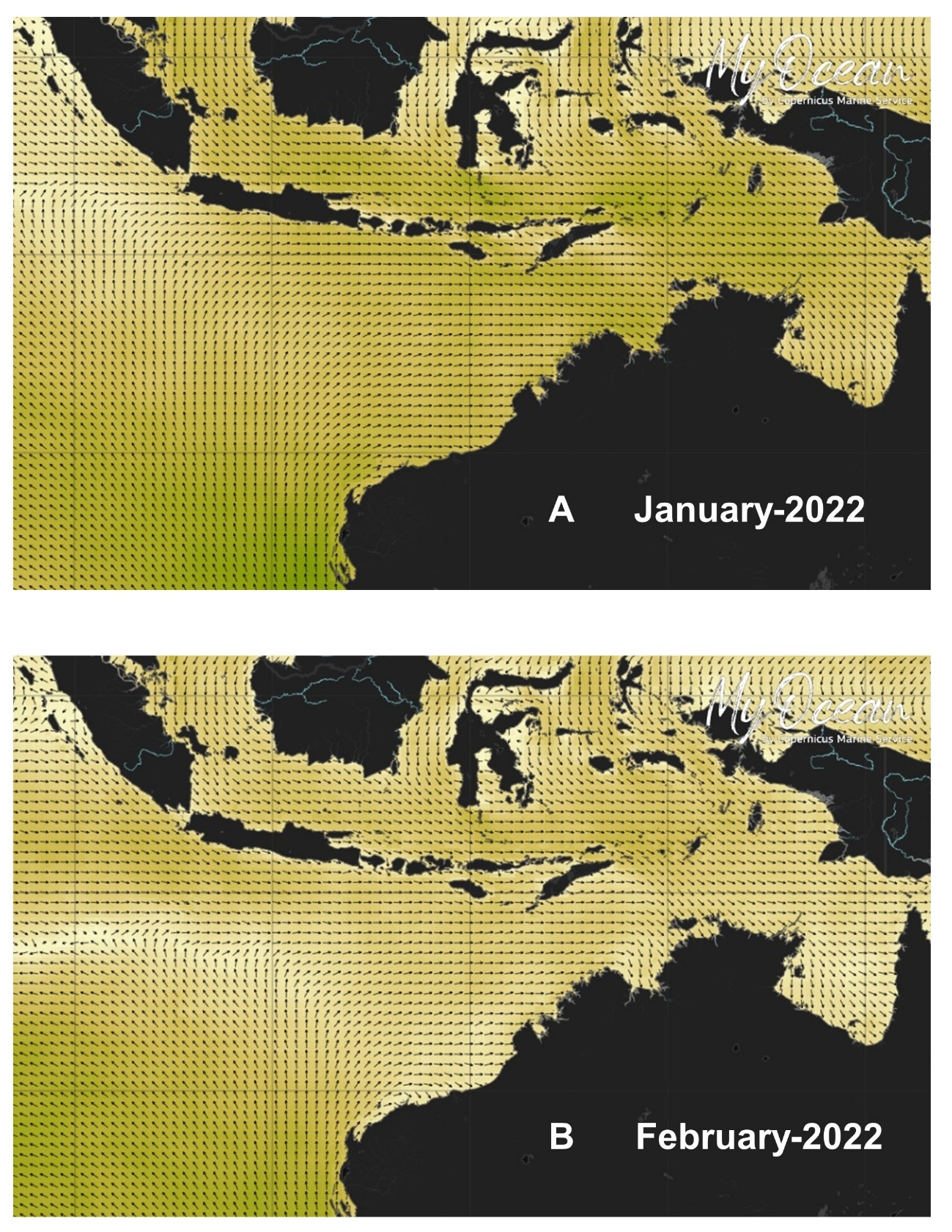
